# Metagenomic alterations in gut microbiota precede and predict onset of colitis in the IL10 gene-deficient murine model

**DOI:** 10.1101/2020.05.04.078022

**Authors:** Jun Miyoshi, Sonny T. M. Lee, Megan Kennedy, Mora Puertolas, Mary Frith, Jason C. Koval, Sawako Miyoshi, Dionysios A. Antonopoulos, Vanessa Leone, Eugene B. Chang

## Abstract

**Background & Aims:** Inflammatory bowel diseases (IBD) are chronic inflammatory disorders where predictive biomarkers for the disease development and clinical course are sorely needed for development of prevention and early intervention strategies that can be implemented to improve clinical outcomes. Since gut microbiome alterations can reflect and/or contribute to impending host health changes, we examined whether gut microbiota metagenomic profiles would provide more robust measures for predicting disease outcomes in colitis-prone hosts.

**Methods:** Using the IL-10 gene-deficient (IL-10 KO) murine model where early life dysbiosis from antibiotic (cefoperozone, CPZ) treated dams vertically-transferred to pups increases risk for colitis later in life, we investigated temporal metagenomic profiles in the gut microbiota of post-weaning offspring and determined their relationship to eventual clinical outcomes.

**Results:** Compared to controls, offspring acquiring maternal CPZ-induced dysbiosis exhibited a restructuring of intestinal microbial membership both in bacteriome and mycobiome that were associated with alterations in specific functional subsystems. Furthermore, among IL-10 KO offspring from CPZ-treated dams, several functional subsystems, particularly nitrogen metabolism, diverged between mice that developed spontaneous colitis (CPZ-colitis) versus those that did not (CPZ-no-colitis) at a time point prior to eventual clinical outcome.

**Conclusions:** Our findings provide support that functional metagenomic profiling of gut microbes has potential and promise meriting further study for development of tools to assess risk and manage human IBD.

**Synopsis:** Currently, predictive markers for the development and course of inflammatory bowel diseases (IBD) are not available. This study supports the notion that gut microbiome metagenomic profiles could be developed into a useful tool to assess risk and manage human IBD.

## Introduction

Inflammatory bowel diseases (IBD) are complex immune disorders that arise from convergence of environmental, microbial, and host genetic factors. Over 200 single-nucleotide polymorphisms identified through genome wide association studies are associated with increased risk for IBD development[1-3], but few, if any, have been clinically useful to predict disease onset or outcomes. Environmental risk factors including diet, hygiene and lifestyle are also important contributors in individuals with a background of IBD genetic susceptibility[4], but even then, no single or combination of factors have emerged as strong disease predictors. The gut microbiome, on the other hand, is highly sensitive to changes in environment and to states of host immune activation which, in theory, could be highly informative in assessing health or impending disease. In this regard, numerous studies have shown both membership and functional changes in the gut microbiome of patients with active IBD, characterized by overall decrease in species diversity, reduced Firmicutes, and increased Proteobacteria compared to non-IBD controls.[5] However, almost without exception, these data have been acquired from samples obtained after disease onset, where immune activation can independently cause dysbiosis. Regardless of whether these changes are cause or consequence of active disease, addressing the unmet need to find high performance predictors of disease onset or relapse in IBD remains a challenge. If achieved, subjects stratified as “high risk” would benefit from early interventions.

Most human IBD microbiome studies have employed marker genes, such as the 16S ribosomal RNA gene, where bacterial membership can be easily profiled. However, these data provide no functional information, which is likely more important in understanding states of host-microbiome interactions. Here, we examine the utility of metagenomic analysis to provide more robust information about the gut microbiome functional state and membership. To do these studies, we used a murine model of vertically transmitted antibiotic (cefaperozone, CPZ)-induced dysbiosis from dams to offspring using IL-10 gene-deficient (IL-10 KO) mice where the pups are prone to develop spontaneous colitis.[6] Like human IBD, disease penetrance in IL-10 KO mice is low, but can be increased when mice are subjected to certain factors such as high fat, low fiber Western-type diet[7], pathobionts (*H. hepaticus*), and other environment shifts that cause gut microbiota perturbations. Thus, this model is sufficiently reflective of human IBD, but also allows for temporal tracking of pre-disease dynamics in both host and microbiome, allowing us to identify potential predictive biomarkers for disease outcomes. We show vertically transmitted maternal peripartum CPZ-induced dysbiosis into genetically susceptible offspring induces shifts in ontology pathways mapped by KEGG that precede and predict which mice go on to develop colitis i.e., CPZ-colitis vs. CPZ-no-colitis. Prominent among them were alterations in pathways associated with microbial nitrogen metabolism, corroborating findings of other groups that had also identified these pathways as being associated with active IBD.[8-11] Also, offspring with vertically transmitted CPZ-induced dysbiosis showed persistent differences in functional profiles as well as microbial composition compared to control offspring of non-CPZ-treated dams. Together, our findings provide proof-of-concept that more robust measures of the gut microbiome, in this case metagenomic profiling, can be the harbingers of host health vs. disease states. We believe the principles identified in our murine colitis model can be extrapolated to the management of IBD in humans. Further studies in humans are needed to delineate if nitrogen metabolism can be used in a similar manner as a marker of dysbiosis. However, our study provides a framework from which to explore this concept and sets the stage for using gut microbiome metagenomics to identify disease markers in IBD patients that could predict relapse of disease that precedes clinical symptoms, unlike the markers of disease currently available, which indicate established relapse and are associated with active symptoms.

## Results

### Effects of maternal peripartum antibiotic exposure on gut microbiome composition of offspring

DNA were extracted and shotgun sequencing was performed on fecal samples harvested from our previously published study. The peripartum CPZ treatment protocol employed is shown in Figure 1. We observed that the incidence of spontaneous colitis among pups from the no-treatment (NT) group was 0% (0/18) in females and 4.3% (1/23) in males, while the incidence in pups from dams exposed to peripartum CPZ (CPZ group) increased to 12.5% (2/16) in females and 30.8% (8/26) in males, respectively. Among the CPZ group mice that developed colitis late in life (CPZ-colitis mice), the mean age of onset was 16.5 weeks of age. MG-RAST[12] was used to assign microbial identity and functions to reads from 51 metagenome samples, representing 3 time points (3, 7, and 11 weeks of age) from representative animals, including 6 NT mice, 5 CPZ-colitis mice, and 6 CPZ-no-colitis mice (mice in CPZ group that did not develop overt colitis). Taxonomic compositional profiles including all bacterial and fungal community members were used to calculate Bray-Curtis dissimilarity, and principal coordinate analysis (PCoA) using permutational multivariate analysis of variance (PERMANOVA) revealed significantly distinct clustering of NT and CPZ groups at 3, 7, and 11 weeks of age (Figure 2A). When broken down into separate bacterial and fungal community analyses, this clustering was recapitulated by both communities across all time points (Supplementary Figure 1).

**Figure 1.**
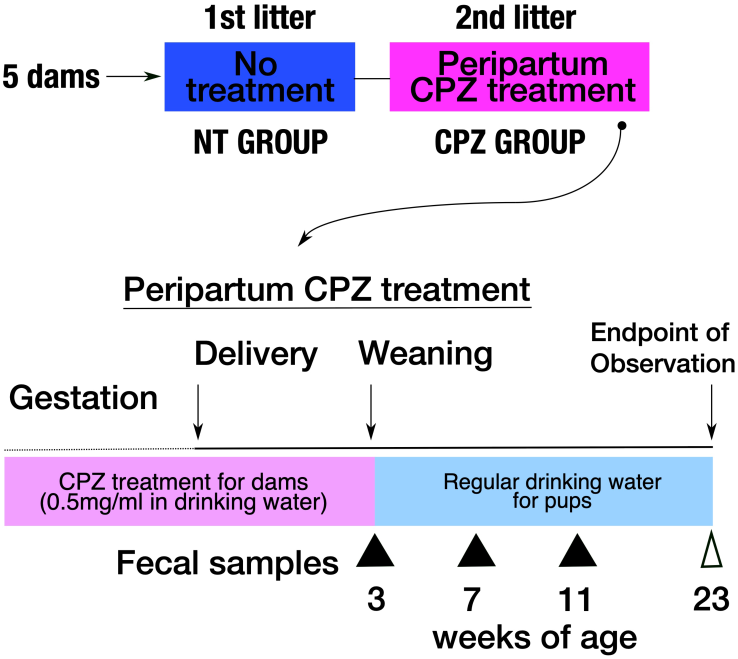
Maternal peripartum CPZ treatment. The first and second litters from five dams were tracked as NT controls (no-treatment) and CPZ (antibiotics treatment) group, respectively. Cefoperazone (CPZ) was administered in dam’s drinking water (0.5 mg/mL) beginning at the third week of the second gestation until weaning of pups (3 weeks of age of pups). After weaning, all pups were provided regular drinking water.

**Figure 2.**
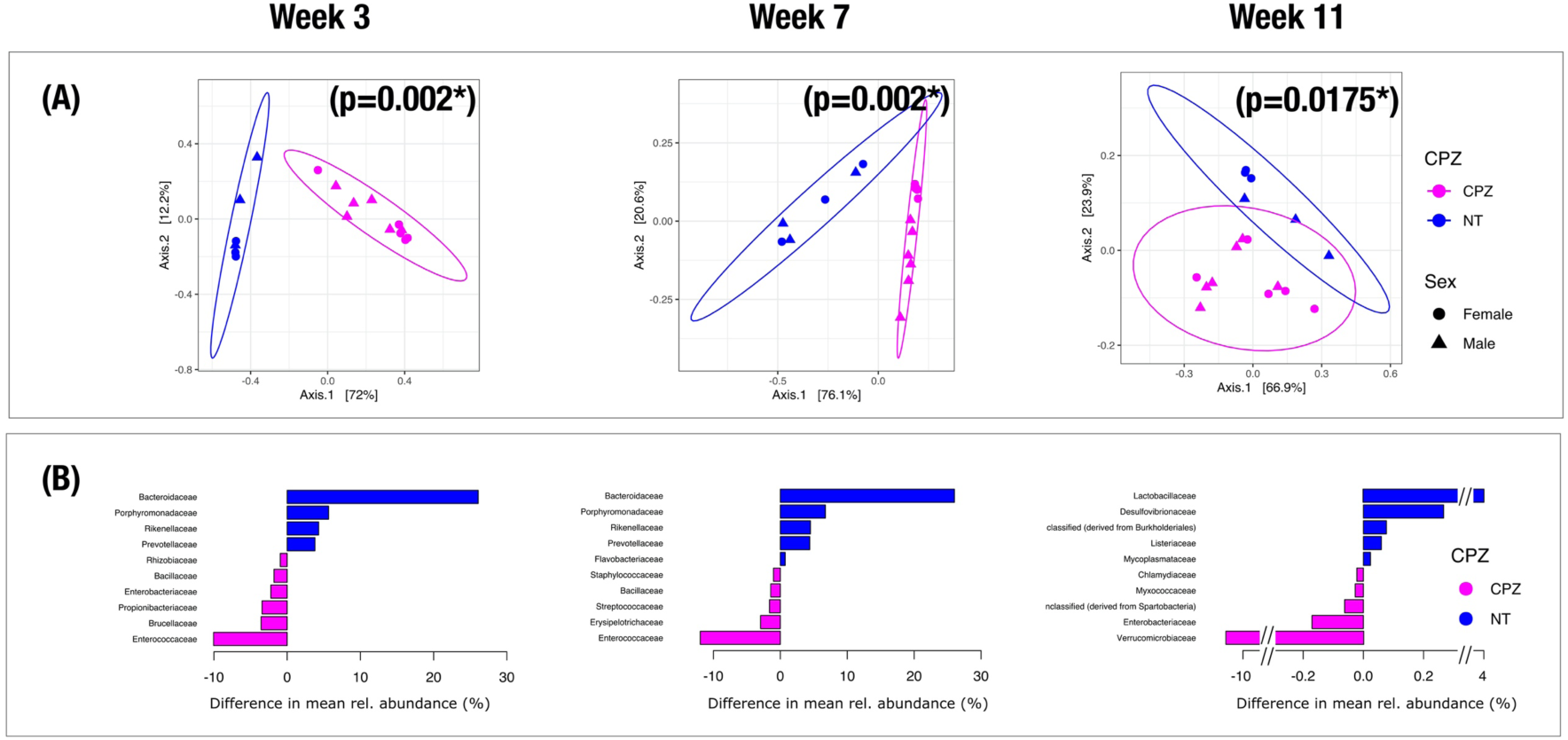
Microbial composition between litters with/without maternal peripartum CPZ exposure. Metagenomic shotgun sequencing data were assigned taxonomy with MG-RAST and analyzed by Bray-Curtis dissimilarity across groups. (A) PCoA plots show that offspring from cefoperazone (CPZ)-treated dams (purple) and no-treatment (NT) controls (blue) formed significantly distinct clusters at all time points. Male mice are represented by triangles, female mice by circles. (B) Family-level relative abundances between CPZ and NT samples were compared and the top 10 differential mean abundance between the two treatment groups are presented.

Family-level relative abundances between CPZ and NT samples were significantly different for 86 bacterial taxa at week 3, and 101 taxa were different at week 7 (Figure 2B, Supplementary Table S1). Among these taxa, *Bacteroidaceae* exhibited the largest difference in relative abundance across treatment groups (week 3: *p* = 0.019; week 7: *p* = 0.007), with enrichment in relative abundance in the NT group (week 3: 33.06 ± 11.94%; week 7: 29.11 ± 9.22%) compared to the CPZ group (week 3: 6.97 ± 4.08%; week 7: 3.09 ± 1.05%). *Enterococcaceae* were also highly differentially abundant (week 3: *p* = 0.019; week 7: *p* < 0.01), but exhibited enrichment in CPZ mice (week 3: 12.56 ± 8.35%; week 7: 14.54 ± 3.57%) relative to NT mice (week 3: 2.50 ± 0.89%; week 7: 2.58 ± 0.67%). At week 11, CPZ and NT mice showed 54 significantly differentially abundant bacterial taxa, with *Verrucomicrobiaceae* exhibiting the largest differences (*p* < 0.01) between CPZ (11.30 ± 2.77%) and NT mice (0.66 ± 0.72%) (Figure 2B, Supplementary Table S1).

No significant differences in gut bacterial community membership were observed between the CPZ-colitis and CPZ-no-colitis groups at weeks 3 or 7 (Figure 3A). However, at week 11, distinct and significant clustering became apparent (*p* = 0.028). These differences were mainly attributed to the presence and abundance of the families *Lactobacillaceae* (colitis: 0.76 ± 0.52%, no-colitis: 3.39 ± 2.64%) and *Enterococcaceae* (colitis: 1.09 ± 0.39%, no-colitis: 2.27 ± 0.76%), which exhibited the largest differences in relative abundances (Figure 3B, Supplementary Table S1).

**Figure 3.**
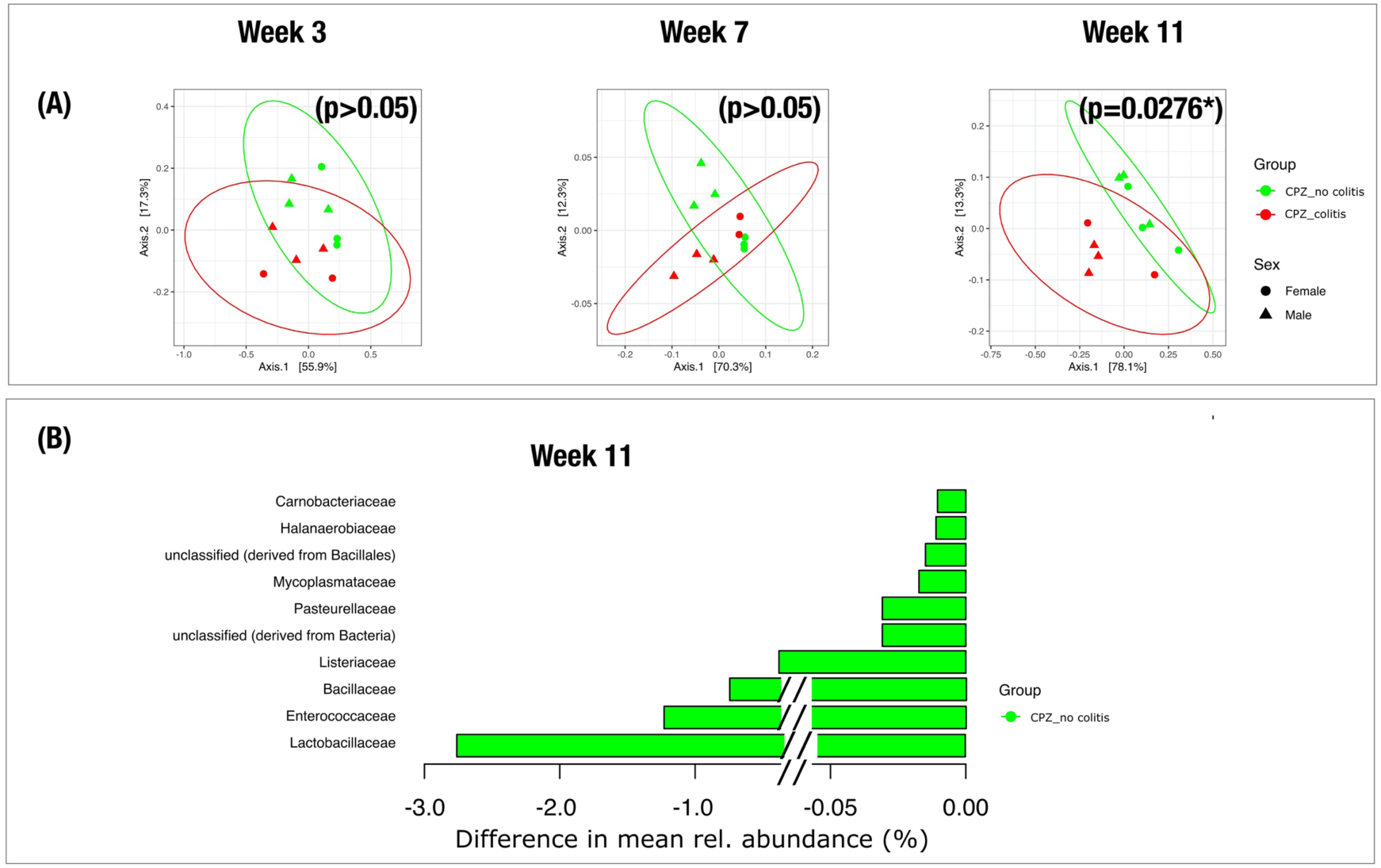
Microbial composition between pups that developed spontaneous colitis later in life and those that did not among maternal peripartum CPZ treated offspring. Metagenomic shotgun sequencing data were assigned taxonomy with MG-RAST and analyzed by Bray-Curtis dissimilarity across groups. (A) Among the CPZ group with maternal peripartum CPZ treatment, pups that developed colitis later in life (red; CPZ-colitis group) or did not develop colitis (green; CPZ-no-colitis group) clustered separately only at week 11. Male mice are represented by triangles, female mice by circles. (B) Family-level relative abundances between CPZ-colitis and CPZ-no-colitis samples at week 11 were compared. The top 10 differential mean abundance between the two resultant colitis phenotypes are presented.

### Maternal peripartum antibiotic exposure impacts offspring gut microbiota functional gene dynamics that correlate with disease course

In addition to significant changes in microbial community membership, we observed persistent functional dysbiosis in pups from CPZ-treated dams relative to those from NT dams. Community-wide functional representation was assessed by mapping metagenomic reads to KEGG Level 3 pathways. PCoA plots based on Bray-Curtis dissimilarity and PERMANOVA clustering analysis revealed significant, distinct clustering of the CPZ and NT groups at all time points (Figure 4A). At weeks 3, 7, and 11, genes related to aminoacyl-tRNA biosynthesis (KEGG pathway ko00970) were overrepresented in NT mice compared to CPZ mice, while at weeks 3 and 7, genes related to ABC transporters (KEGG pathway ko02010) were enriched in CPZ-treated mice compared to NT mice (Figure 4B, Supplementary Table S2).

**Figure 4.**
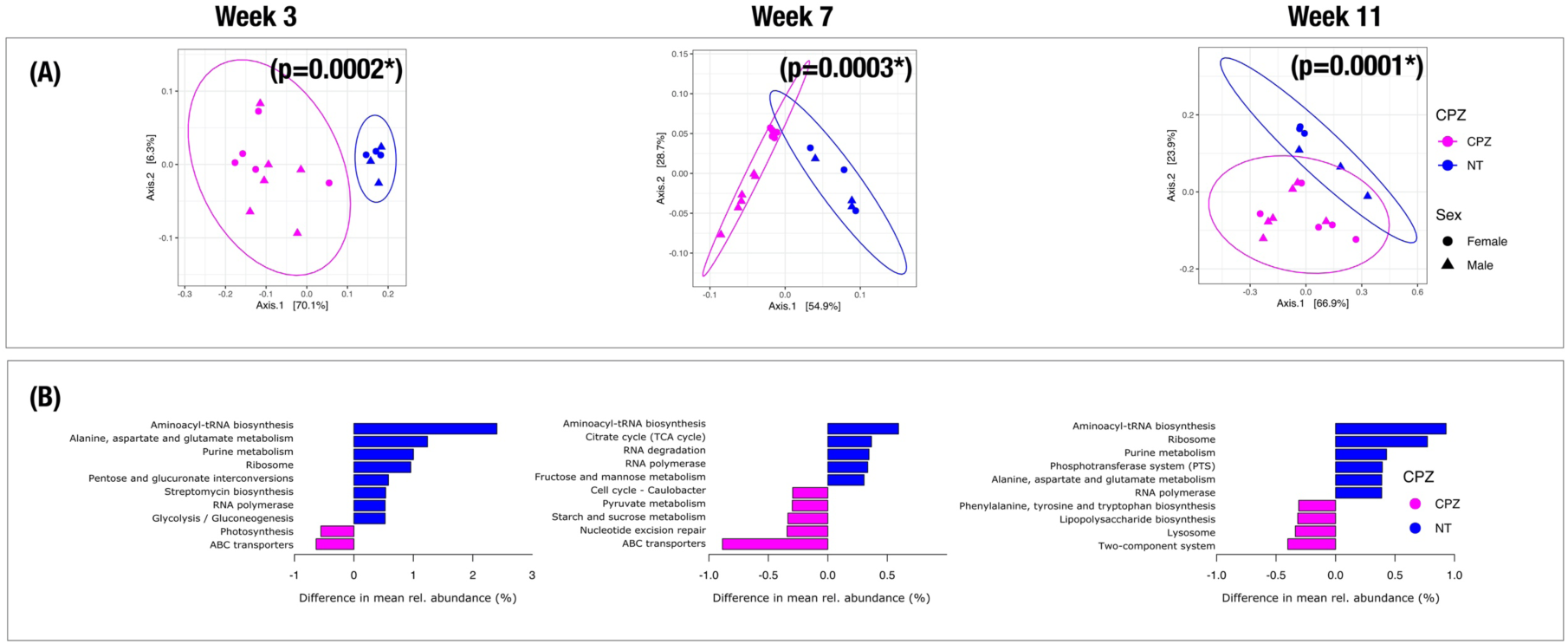
Functional pathways between litters with/without maternal peripartum CPZ exposure. Metagenomic shotgun sequencing data were mapped to KEGG Level 3 functional pathways with MG-RAST and analyzed by Bray-Curtis dissimilarity across treatment groups. (A) PCoA plots of community-wide functional profile show that offspring from cefoperazone (CPZ)-treated dams (purple) and no-treatment (NT) controls (blue) formed significantly distinct clusters at all time points. Male mice are represented by triangles, female mice by circles. (B) The top 10 differential mean abundance gene functional pathways across the CPZ-treated group (purple) and NT group (blue) at each time point are presented.

We next assessed whether functional pathway representation changed over time for either treatment group (Figure 5A). PCoA and clustering analysis revealed that for NT mice, the functional profile did not change significantly across time points (Figure 5B), whereas for CPZ mice, functional profiles formed significantly distinct clusters at each time point (Figure 5C, *p* < 0.01). Interestingly, at week 11, the CPZ-no-colitis group appeared to cluster between the CPZ-colitis group and the NT group (Figure 5A).

**Figure 5.**
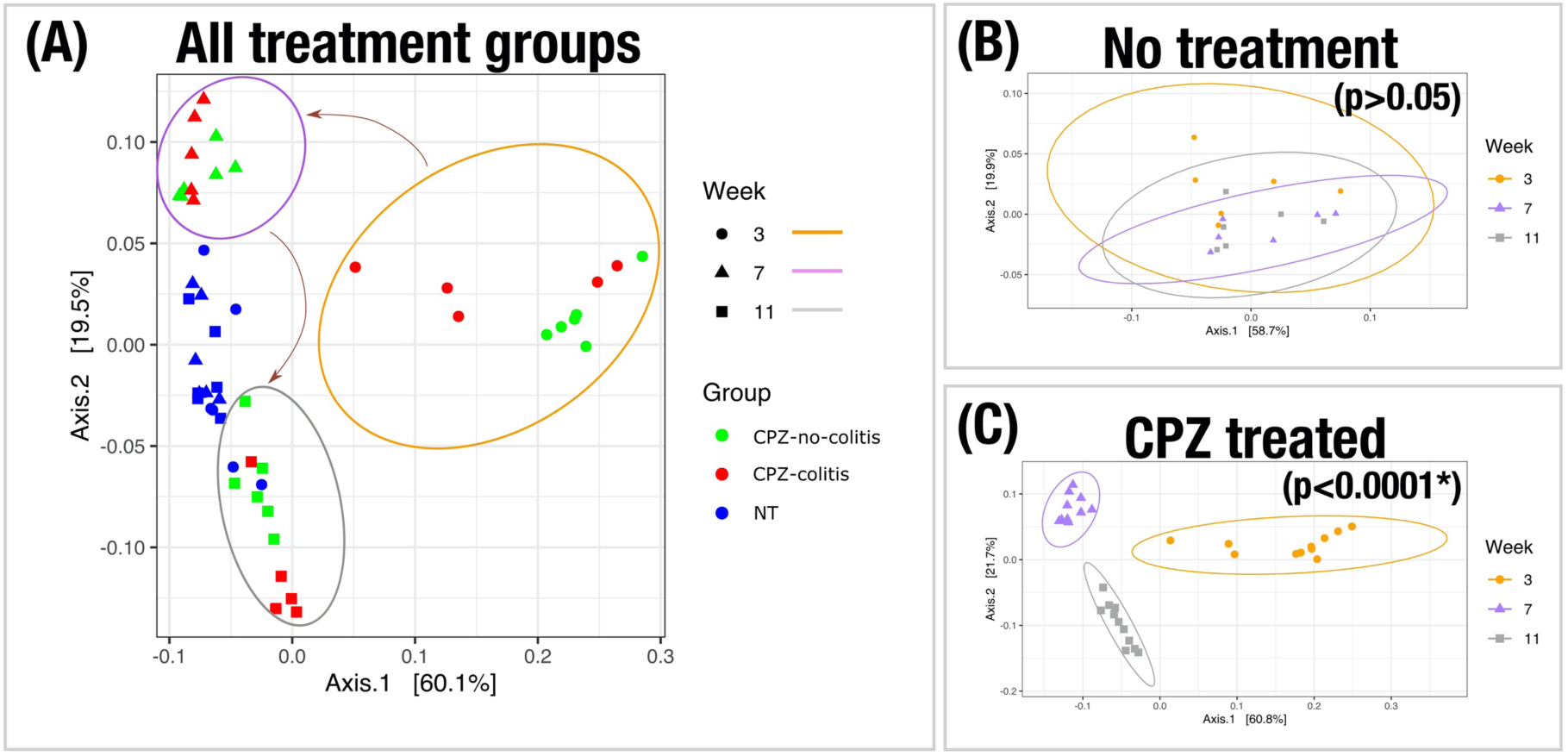
Functional pathway profiles of offspring with maternal peripartum CPZ treatment shifts over time. Metagenomic shotgun sequencing data were mapped to KEGG Level 3 functional pathways, analyzed by Bray-Curtis dissimilarity across time points and ordinated by PCoA. (A) No-treatment (NT) control mice (blue), pups in the cefoperazone (CPZ)-treated group that developed colitis later in life (red; CPZ-colitis group), and pups in the CPZ-treated group that did not develop colitis (green; CPZ-no-colitis group) are plotted together. Weeks are denoted by shape; among the CPZ-treated group, weeks are further denoted by colored ellipses. Arrows indicate transitions of CPZ-treated mice over time. (B) NT control mice showed no significant differences in clustering across time points. (C) CPZ-treated mice clustered distinctly at each time point.

As with gut microbial taxonomic composition, we observed significant differences in functional pathway representation between CPZ-colitis and CPZ-no-colitis mice at week 11 but not weeks 3 or 7 (KEGG pathway level 3) (Figure 6A). At week 11, while none of the mice had yet developed frank symptoms of spontaneous colitis, genes related to nitrogen metabolism (KEGG pathway ko00910) and protein digestion and absorption (KEGG pathway ko04974) were significantly differentially abundant between the mice that would eventually develop colitis, and those that would not (Figure 6B, Supplementary Table S2). Genes from these pathways were both enriched in the CPZ-colitis group (nitrogen metabolism: 0.009 ± 0.001%; protein digestion and absorption: 0.032 ± 0.002%) as compared to the CPZ-no-colitis group (nitrogen metabolism: 0.001 ± 0.001%; protein digestion: 5.41 × 10^−5^ ± 5.11 × 10^−5^%).

**Figure 6.**
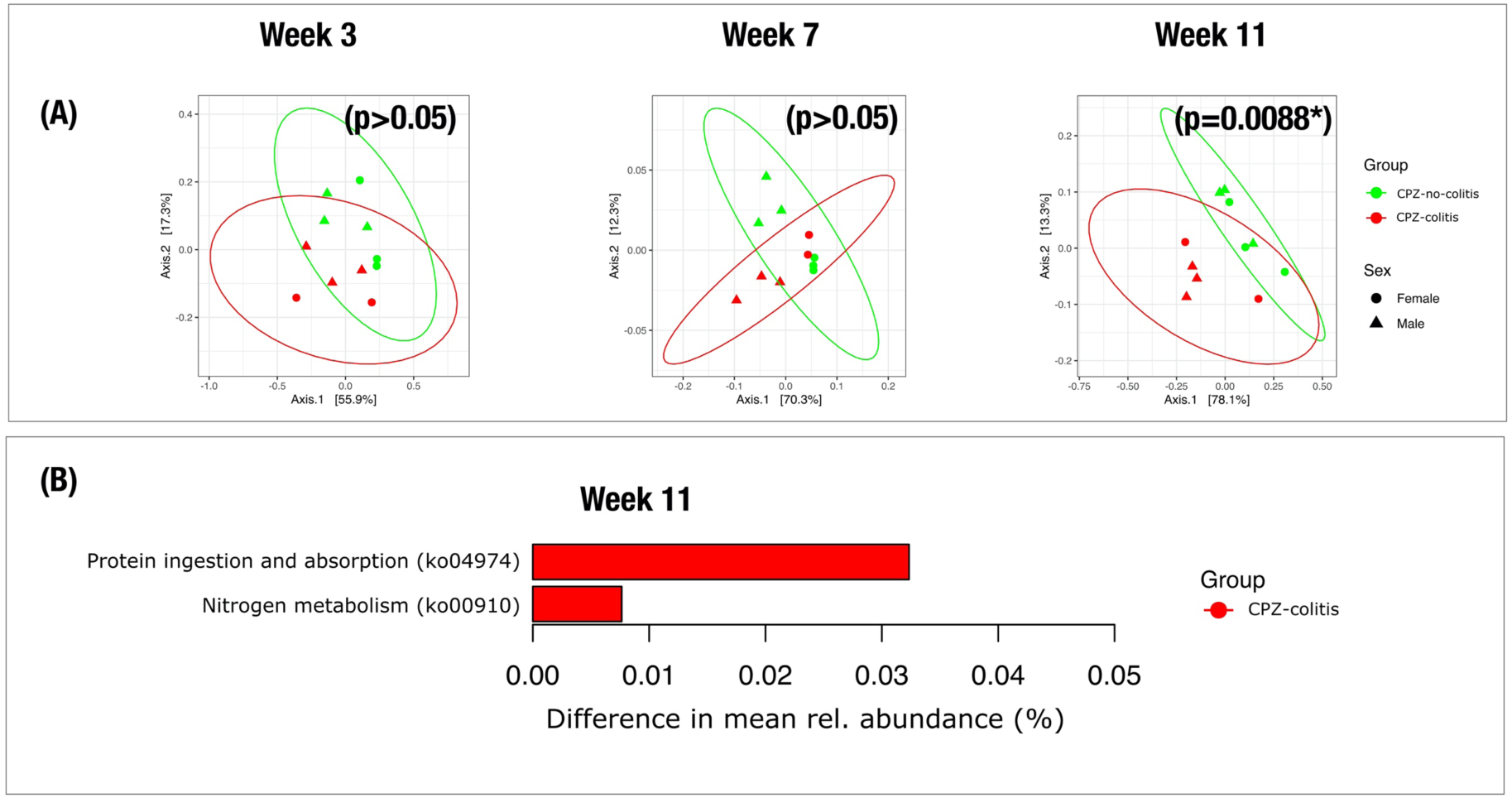
Functional pathways between pups that developed spontaneous colitis later in life and those that did not among offspring with maternal peripartum CPZ treatment. Metagenomic shotgun sequencing data were mapped to KEGG Level 3 functional pathways with MG-RAST and analyzed by Bray-Curtis dissimilarity across treatment groups. (A) Among offspring from cefoperazone (CPZ)-treated dams, pups that developed colitis later in life (red; CPZ-colitis group) and those that did not (green; CPZ-no-colitis group) formed significantly distinct clusters only at week 11. Male mice are represented by triangles, female mice by circles. (B) Bar plots display significantly different functional pathways at weeks 7 and 11 between the CPZ-colitis group (red) vs. the CPZ-no-colitis group (green).

## Discussion

Currently no reliable clinical indicators of impending colitis development or relapse exist, and clinical outcomes in IBD remain unpredictable among genetically susceptible individuals. Advances in this area are hindered by the lack of longitudinal data preceding colitis onset mainly because the spontaneous nature of human IBD largely precludes such studies. The lack of indicators limits the ability of monitor treatment response and to implement preventative measures in the clinic. Since the gut microbiota is strongly implicated in IBD pathogenesis and clinical stool sampling is routine and non-invasive, identifying microbial signatures or metabolites that reliably predict disease onset or relapse would be a major advance in the clinical management of IBD. Here, to explore microbiota-based indicators of future IBD onset or relapse, we analyzed longitudinal changes in gut microbial metagenomes of colitis-prone mice raised in the same environment, some of which developed colitis, while others did not. Importantly, we found altered microbial signatures (particularly those involved in nitrogen metabolism), beginning between 7-11 weeks of age, defined the gut microbiota of mice that later developed colitis, although none of the mice showed clinical symptoms at these time points. Our findings highlight the potential for using alterations in microbial community function to warn of colitis onset and/or relapse in high risk individuals, facilitating timely precautionary measures or prompt intervention.

The present study was prompted by our previous observations that among genetically identical colitis-prone IL-10 KO pups that acquire maternal antibiotic-induced dysbiosis through vertical transmission (CPZ group), about a third developed spontaneous colitis (CPZ-colitis mice) while the others did not (CPZ-no-colitis mice) in contrast to almost no spontaneous colitis development in the NT group.[6] This allowed us to investigate microbial predictors of colitis onset among susceptible pups in this model. We hypothesized that among susceptible individuals raised in the same environment, a divergence in the functional profile of the gut microbiota would precede colitis onset. Using fecal samples collected from these mice at 3, 7, 11 weeks of age, our metagenomic shotgun sequencing phylogenetic and functional analyses revealed that although we observed no differences in gut microbial composition and function between CPZ-colitis and CPZ-no-colitis mice at 3 or 7 weeks of age, the microbiota of the two groups diverged at 11 weeks of age, with the CPZ-no-colitis group becoming more similar to the NT control group, and the CPZ-colitis group diverging in a different direction. This divergence reflected alterations in several gene ontology KEGG pathways, but nitrogen metabolism, as indicated by the differing phylogenetic compositions and functional pathways enriched in the fecal microbiota between groups, was particularly prominent and has been observed by others in active IBD.[10] In spite of this shift, none of the animals showed clinical symptoms of colitis at 11 weeks of age, such as body weight loss, loose stool, or ruffled fur[6]. Thus, this compositional and functional divergence of the gut microbiota preceded and predicted the colitis development.

Several previous studies support the notion that microbial nitrogen metabolism might be associated with the development of overt colitis. Metagenomic shotgun sequencing of fecal DNA samples from pediatric CD patients revealed pathways involved in nitrogen metabolism were significantly more abundant in CD versus control samples.[9]. Additionally, bacterial nitrogen flux via urease activity in mice was associated with colitis severity[10] and microbial ammonia biosynthesis pathways were more highly expressed in a mouse model of IBD compared to controls.[11] Winter et al. demonstrated that nitrate-respiration, through the reduction of host-derived nitrate, provides a growth benefit to facultative anaerobes, such as Enterobacteriaceae, in the inflamed intestine.[8]

While different functional profiles between CPZ-colitis and -no-colitis mice are thought to be a reflection of multiple aspects of altered nitrogen metabolism, fundamentally, they comprised a distinct signature in these mice that predicted impending colitis onset. Indeed, a single universal indicator of impending colitis may not exist; rather, our findings may instead implicate a functional divergence of the gut microbiota from a previous steady state as the most reliable indicator of impending colitis onset or relapse. In a clinical context, longitudinal stool sampling of patients in remission until flare could help to identify patient-specific divergent factors that indicate risk for relapse. In particular, given the high rates of colitis recurrence in UC patients after ileal pouch-anal anastomosis[13] and in CD patients after bowel resection[14], these are clinical models in whom prospective sampling could reveal microbiome-based signatures that predict colitis onset.

Another notable point is that 16S rRNA gene amplicon sequencing, which is standard in microbiome research and was utilized in our previous study[6], could not resolve important differences between groups that were evident from metagenomic shotgun sequencing. Relying solely on 16S rRNA amplicon sequencing data would have been misleading. Metagenomes provide much more robust information about microbial communities and enable phenotypic predictions with greater accuracy and precision [15] and should be considered in future studies, particularly in the context of IBD. While routine metagenomic sequencing of patient samples is currently not practical, studies that utilize metagenomic sequencing and functional profiling may facilitate the development of cost effective, targeted assays to assess patient risk for relapse.

We do concede several limitations of our study. First, it does not test microbially-mediated mechanisms involved in colitis development, and further studies are needed to flesh out our finding that altered nitrogen metabolism preceded the onset of colitis and identify whether this is cause or effect. Nonetheless, our findings provide “proof of concept” that metagenome-derived information can be predictive of risk for disease, in this case the development of colitis associated with antibiotic-induced early life dysbiosis and loss of immune tolerance. These factors themselves were not sufficient to cause disease but set the stage for other factors that can tip the balance in a genetically susceptible host. Second, the murine and human microbiome, environment, diet, and genetics are not equivalent and thus the microbial signatures that predicted colitis in this model may not represent predictive signatures in humans. However, using a murine model enabled the collection of longitudinal data from birth to colitis onset in adulthood, which is unfeasible if not impossible to collect in humans. This model is therefore helpful in developing working paradigms that are relevant to human IBD. The specific metagenomic markers, mapping, and ontology pathways that will be useful to human IBD, however, cannot rely on those found in mouse models and will have to be determined through clinical investigations done prospectively.

Notwithstanding, this study has significant implications for the care of IBD patients, particularly those in remission, who, given the nature of IBD, are at high risk for relapse. The state and changes of the gut microbiome are very sensitive to what is happening in the host; in turn, these changes can feed forward in contributing to the underlying problem or perturbation. We propose that the microbiome is, at minimum, the “canary in the coal mine” to alert physicians about states of normalcy or impending changes. These changes are quantifiable by clustering patterns or functional subsystems that can provide metrics for clinical state and course and can potentially provide insight into cause and/or effect relationships. Irrespective of cause and effect, however, the crucial point from this study is that metagenomic states and changes may serve as indicators of what is happening to the host. Such indicators would be clinically useful for introducing early interventions in the management of IBD. For example, if microbial community function of an individual in remission begins to diverge, this may be a sign of impending colitis— longitudinal sampling is needed to make this judgement call. Additionally, serial sampling could provide an opportunity to steer a patient’s diverging microbiome back to steady state by supplementation with the patient’s own remission microbiota samples.

In conclusion, the findings of the present study strongly suggest that functional divergence of the gut microbiota, potentially in the form of microbial nitrogen metabolism, warrants further study as a potential indicator for IBD risk or relapse. The most practical application of such research in the clinical setting might be for monitoring patients in remission. One goal should be to develop a scalable and cost-effective assay to guide the implementation of preventative measures and to improve clinical management and monitoring of patient treatment response. The current lack of biomarkers and other metrics to guide clinical management of IBD highlight the critical need for further investigation into promising microbiome-based metrics.

## Methods

### Fecal DNA Samples for Metagenomic Shotgun Sequencing

DNA were extracted from feces of *H. hepaticus*-free SPF IL-10 KO mice with a C57BL/6J genetic background, that were harvested in our previous study.[6] Briefly, five breeding pairs were prepared after the normalization of microbiota by bedding transfer[16] to obtain offspring of non-treatment (NT) group and cefoperazone (CPZ)-treatment group. The first and second litters from the five breeding pairs were tracked as NT controls and CPZ group, respectively. The peripartum CPZ treatment consisted of CPZ (0.5 mg/mL) administration in the dam’s drinking water beginning at the third week of the second gestation until the weaning of pups (3 weeks of age of pups) as shown in Figure 1. The pups were tracked until 23 weeks of age. Collected feces were rapidly frozen at −80°C and DNA was extracted as previously described.[17] NT litters did not develop overt spontaneous colitis throughout the observation period until 23 weeks of age, with the exception of one male, while the CPZ group showed a higher incidence of overt spontaneous colitis, i.e. 2 out of 16 females and 8 out of 26 males developed colitis (the mean age of onset was 16.5 weeks of age). Given this observation, fecal DNA samples described below were selected for metagenomic shotgun sequencing analysis. As representative samples of NT offspring, fecal DNA samples at 3, 7, and 11 weeks of age obtained from 3 females and 3 males in the NT group that did not develop frank, spontaneous colitis during the observation period were analyzed. For the CPZ group, fecal DNA samples at 3, 7, and 11 weeks of age from 3 females and 3 males that did not develop overt spontaneous colitis during the observation period were examined as the CPZ-no-colitis group. Meanwhile, the time-series samples from 2 females and 3 males that developed overt colitis later in life, i.e. these mice did not show frank colitis symptoms at 11 weeks of age, were investigated as CPZ-colitis group.

### Metagenomic analyses

Metagenome libraries were generated using the Illumina Hiseq platform. An average of 16,438,078 sequences per sample were generated in this study (Supplementary Table S1). We used the quality control filter, an internal tool in the MG-RAST v4.0.3 pipeline[12] and excluded an average of 764,372,216 reads from further analyses (Supplementary Table S3). Of these, dereplication identified an average of 5% as clusters of artificially replicated sequences. The filter parameters included a cutoff value of 0.9, with no length difference requirement and an initial base pair match of 3 base pairs. After quality filtering, an average of 1,405,454,114 sequences for the metagenomes were used in the metagenomic analyses (Supplementary Table S3). Approximately an average of 50% of our reads were annotated (e-value cutoff of 1e^-05^) to an assigned function or specific gene by MG-RAST v4.0.3 pipeline (Supplementary Table S4), against the SEED database.

### Statistics

Principal coordinate analysis (PCoA) plots were generated based on Bray-Curtis dissimilarity using the R (https://www.R-project.org/) package phyloseq[18] and clustering analysis was performed by permutational multivariate analysis of variance (PERMANOVA) using the R package vegan[19]. Statistical differences between metagenome profiles were calculated based on the Fisher’s exact test with corrected q-values (Storey’s FDR multiple test correction approach) using the software package STAMP v.2.1.3.[20] The criterion for statistical significance was set at *p* < 0.05 or FDR < 0.05.

## Abbreviations

IBD: inflammatory bowel diseases,
UC: ulcerative colitis,
CD: Crohn’s disease,
CPZ: cefoperazone,
BNF: biological nitrogen fixation,
PCoA: principal coordinates analysis,
(PERMANOVA): permutational multivariate analysis of variance

## Data Availability

The raw metagenomic sequencing data in this study is publicly available at MG-RAST under the accession number mgp82768 and at the NCBI Sequence Read Archive under the accession number SRP252264.

## Access to Data

All authors had access to the entire dataset and have reviewed and approved the final manuscript.

## References

[1] Liu JZ, van Sommeren S, Huang H, Ng SC, Alberts R, Takahashi A, Ripke S, Lee JC, Jostins L, Shah T, Abedian S, Cheon JH, Cho J, Dayani NE, Franke L, Fuyuno Y, Hart A, Juyal RC, Juyal G, Kim WH, Morris AP, Poustchi H, Newman WG, Midha V, Orchard TR, Vahedi H, Sood A, Sung JY, Malekzadeh R, Westra HJ, Yamazaki K, Yang SK, International Multiple Sclerosis Genetics C, International IBDGC, Barrett JC, Alizadeh BZ, Parkes M, Bk T, Daly MJ, Kubo M, Anderson CA, Weersma RK. Association analyses identify 38 susceptibility loci for inflammatory bowel disease and highlight shared genetic risk across populations. Nat Genet 2015;47(9):979–86.

[2] Liu TC, Stappenbeck TS. Genetics and Pathogenesis of Inflammatory Bowel Disease. Annu Rev Pathol 2016;11:127–48.

[3] Brant SR, Okou DT, Simpson CL, Cutler DJ, Haritunians T, Bradfield JP, Chopra P, Prince J, Begum F, Kumar A, Huang C, Venkateswaran S, Datta LW, Wei Z, Thomas K, Herrinton LJ, Klapproth JA, Quiros AJ, Seminerio J, Liu Z, Alexander JS, Baldassano RN, Dudley-Brown S, Cross RK, Dassopoulos T, Denson LA, Dhere TA, Dryden GW, Hanson JS, Hou JK, Hussain SZ, Hyams JS, Isaacs KL, Kader H, Kappelman MD, Katz J, Kellermayer R, Kirschner BS, Kuemmerle JF, Kwon JH, Lazarev M, Li E, Mack D, Mannon P, Moulton DE, Newberry RD, Osuntokun BO, Patel AS, Saeed SA, Targan SR, Valentine JF, Wang MH, Zonca M, Rioux JD, Duerr RH, Silverberg MS, Cho JH, Hakonarson H, Zwick ME, McGovern DP, Kugathasan S. Genome-Wide Association Study Identifies African-Specific Susceptibility Loci in African Americans With Inflammatory Bowel Disease. Gastroenterology 2017;152(1):206–17 e2.

[4] Molodecky NA, Soon IS, Rabi DM, Ghali WA, Ferris M, Chernoff G, Benchimol EI, Panaccione R, Ghosh S, Barkema HW, Kaplan GG. Increasing incidence and prevalence of the inflammatory bowel diseases with time, based on systematic review. Gastroenterology 2012;142(1):46-54 e42; quiz e30.

[5] Gevers D, Kugathasan S, Knights D, Kostic AD, Knight R, Xavier RJ. A Microbiome Foundation for the Study of Crohn’s Disease. Cell Host Microbe 2017;21(3):301–4.

[6] Miyoshi J, Bobe AM, Miyoshi S, Huang Y, Hubert N, Delmont TO, Eren AM, Leone V, Chang EB. Peripartum Antibiotics Promote Gut Dysbiosis, Loss of Immune Tolerance, and Inflammatory Bowel Disease in Genetically Prone Offspring. Cell Rep 2017;20(2):491–504.

[7] Devkota S, Wang Y, Musch MW, Leone V, Fehlner-Peach H, Nadimpalli A, Antonopoulos DA, Jabri B, Chang EB. Dietary-fat-induced taurocholic acid promotes pathobiont expansion and colitis in Il10-/-mice. Nature 2012;487(7405):104–8.

[8] Winter SE, Winter MG, Xavier MN, Thiennimitr P, Poon V, Keestra AM, Laughlin RC, Gomez G, Wu J, Lawhon SD, Popova IE, Parikh SJ, Adams LG, Tsolis RM, Stewart VJ, Baumler AJ. Host-derived nitrate boosts growth of E. coli in the inflamed gut. Science 2013;339(6120):708–11.

[9] Lewis JD, Chen EZ, Baldassano RN, Otley AR, Griffiths AM, Lee D, Bittinger K, Bailey A, Friedman ES, Hoffmann C, Albenberg L, Sinha R, Compher C, Gilroy E, Nessel L, Grant A, Chehoud C, Li H, Wu GD, Bushman FD. Inflammation, Antibiotics, and Diet as Environmental Stressors of the Gut Microbiome in Pediatric Crohn’s Disease. Cell Host Microbe 2015;18(4):489–500.

[10] Ni J, Shen TD, Chen EZ, Bittinger K, Bailey A, Roggiani M, Sirota-Madi A, Friedman ES, Chau L, Lin A, Nissim I, Scott J, Lauder A, Hoffmann C, Rivas G, Albenberg L, Baldassano RN, Braun J, Xavier RJ, Clish CB, Yudkoff M, Li H, Goulian M, Bushman FD, Lewis JD, Wu GD. A role for bacterial urease in gut dysbiosis and Crohn’s disease. Sci Transl Med 2017;9(416).

[11] Sharpton T, Lyalina S, Luong J, Pham J, Deal EM, Armour C, Gaulke C, Sanjabi S, Pollard KS. Development of Inflammatory Bowel Disease Is Linked to a Longitudinal Restructuring of the Gut Metagenome in Mice. mSystems 2017;2(5).

[12] Meyer F, Paarmann D, D’Souza M, Olson R, Glass EM, Kubal M, Paczian T, Rodriguez A, Stevens R, Wilke A, Wilkening J, Edwards RA. The metagenomics RAST server - a public resource for the automatic phylogenetic and functional analysis of metagenomes. BMC Bioinformatics 2008;9:386.

[13] Murrell ZA, Melmed GY, Ippoliti A, Vasiliauskas EA, Dubinsky M, Targan SR, Fleshner PR. A prospective evaluation of the long-term outcome of ileal pouch-anal anastomosis in patients with inflammatory bowel disease-unclassified and indeterminate colitis. Dis Colon Rectum 2009;52(5):872–8.

[14] Swoger JM, Regueiro M. Evaluation for postoperative recurrence of Crohn disease. Gastroenterol Clin North Am 2012;41(2):303–14.

[15] Ranjan R, Rani A, Metwally A, McGee HS, Perkins DL. Analysis of the microbiome: Advantages of whole genome shotgun versus 16S amplicon sequencing. Biochem Biophys Res Commun 2016;469(4):967–77.

[16] Miyoshi J, Leone V, Nobutani K, Musch MW, Martinez-Guryn K, Wang Y, Miyoshi S, Bobe AM, Eren AM, Chang EB. Minimizing confounders and increasing data quality in murine models for studies of the gut microbiome. PeerJ 2018;6:e5166.

[17] Wang Y, Hoenig JD, Malin KJ, Qamar S, Petrof EO, Sun J, Antonopoulos DA, Chang EB, Claud EC. 16S rRNA gene-based analysis of fecal microbiota from preterm infants with and without necrotizing enterocolitis. ISME J 2009;3(8):944–54.

[18] McMurdie PJ, Holmes S. phyloseq: an R package for reproducible interactive analysis and graphics of microbiome census data. PLoS One 2013;8(4):e61217.

[19] Oksanen JB, F.G.; Friendly, M.; Kindt, R.; Legendre, P.; McGlinn, D.; Minchin, P.R.; O’Hara, R.B.; Simpson, G.L.; Solymos, P.; Stevens, M.H.H.; Szoecs, E.; Wagner, H. vegan: Community Ecology Package. R package version.5-6.; 2019. Available from: https://CRAN.R-project.org/package=vegan. [Accessed 04/27 2020].

[20] Parks DH, Beiko RG. Identifying biologically relevant differences between metagenomic communities. Bioinformatics 2010;26(6):715–21.

