## Supplementary Figure S1 for "Metagenomic alterations in gut microbiota precede and predict onset of colitis in the IL10 gene-deficient murine model"

### Supplementary Figure 1.

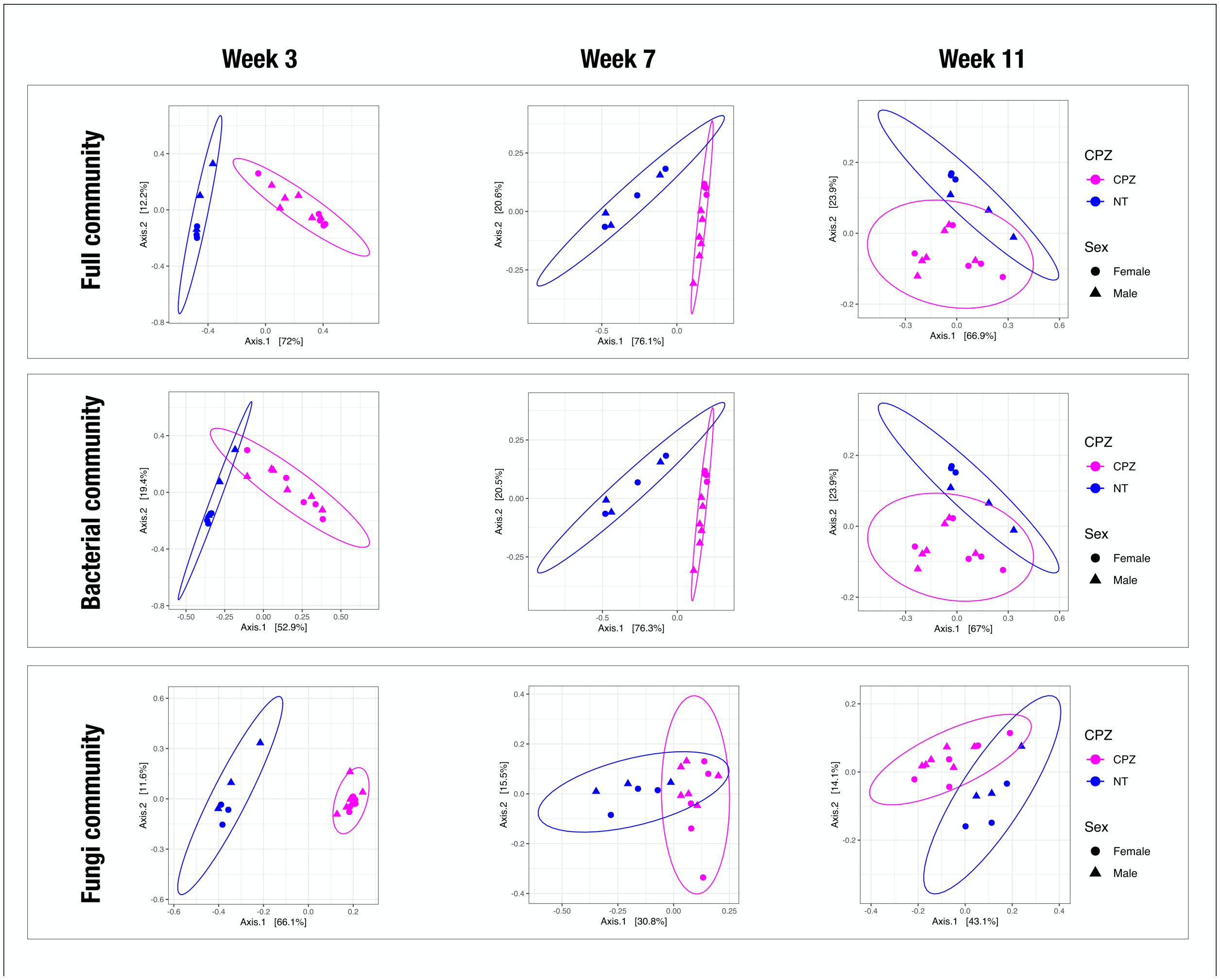


**Supplementary Figure S1. Microbial membership between litters with/without maternal peripartum CPZ exposure.**

Metagenomic shotgun sequencing data were assigned taxonomy with MG-RAST and analyzed by Bray-Curtis dissimilarity across groups. Similar to the overall microbial community membership (Figure 2A), the composition of both microbiome and mycobiome varied between CPZ and NT groups at all time points and is different between pups from cefoperazone (CPZ)-treated dams (purple) and no-treatment (NT) controls (blue), indicated by significantly distinct clusters at all time points.
